# Multiscale RNA editing analysis reveals cell type–specific regulatory programs across disease states in acute myeloid leukemia

**DOI:** 10.64898/2026.02.04.703797

**Authors:** Tongjun Gu, Doan Bui, Guru Subramanian Guru Murthy, Anne E. Kwitek

**Author notes:** Corresponding author: Tongjun Gu, PhD, Versiti Blood Research Institute, 8727 W Watertown Plank Rd, Milwaukee, WI 53226.

## Abstract

**Background:** Acute myeloid leukemia (AML) is characterized by marked cellular heterogeneity and immune dysregulation. Adenosine-to-inosine (A-to-I) RNA editing, primarily catalyzed by ADAR and ADARB1, represents an important post-transcriptional regulatory mechanism, yet its condition- and cell type–specific landscape in AML remains poorly defined, particularly at single-cell resolution.

**Methods:** We analyzed publicly available single-cell RNA sequencing data from healthy donors (HL), newly diagnosed AML (ND), remission (RM), and persistent disease (PO), integrating single-cell and pseudo-bulk analyses in a multiscale framework. RNA editing sites were identified using a stringent discovery pipeline and quantified at both pseudo-bulk and cell type–resolved levels. Differential RNA editing was assessed using regression-based read-count models, primarily beta-binomial regression with subject-specific random effects when applicable. Pairwise contrasts between clinical conditions were evaluated using delta-method inference, with statistical significance defined by false discovery rate and a minimum effect-size threshold. Selected editing sites were examined in independent human AML cohorts for validation and clinical association.

**Results:** We identified 2,875 recurrent A-to-I RNA editing sites enriched in intronic and 3′ untranslated regions and linked to immune and inflammatory pathways. At the pseudo-bulk level, 150 sites were differentially edited across clinical states, and global RNA editing varied by condition, showing an overall negative association with ADAR and ADARB1 expression with context-dependent exceptions. Cell type–resolved analyses identified 148 differentially edited sites with strong lineage specificity. In ND, leukemia-associated cell states consistently exhibited lower editing than lineage-matched healthy counterparts. T cells consistently harbored differential editing signals across all condition contrasts, while progenitor-like cells showed the strongest RM-versus-ND differences despite minimal changes in global editing. Notable editing events were observed in *GBP4, SPN, TNFSF10, EMB*, and *FKBP5*. Several candidate sites were validated in independent AML cohorts and were associated with clinical features.

**Conclusions:** This multiscale analysis reveals that RNA editing in AML is condition- and cell type–specific and is not fully captured by bulk transcriptomic measures. Site-specific, lineage-restricted RNA editing represents a distinct regulatory layer that reflects disease state and cellular context, highlighting its potential relevance for understanding AML biology and informing future biomarker development.

## Introduction

Acute myeloid leukemia (AML) is an aggressive hematologic malignancy characterized by clonal expansion of immature myeloid precursors arising from hematopoietic stem and progenitor cells.[1] Diagnosis, risk stratification, and many treatment decisions are guided by genetic and cytogenetic findings.[2] Although most patients achieve an initial remission with induction chemotherapy, relapse is common, and 5-year overall survival remains ∼30%.[1] These clinical realities underscore the role of non-genetic contributors to functional heterogeneity beyond therapy-resistant genomic clones.

RNA editing is a post-transcriptional mechanism that diversifies the transcriptome by altering RNA sequences. In humans, two major types of RNA editing have been described—adenosine-to-inosine (A-to-I) and cytidine-to-uridine (C-to-U)—with A-to-I editing being the predominant form. A-to-I editing is catalyzed by members of the adenosine deaminase acting on RNA (ADAR) family, primarily *ADAR* and *ADARB1*. Because inosine is read as guanine by the reverse transcription and translation machinery, A-to-I editing is often detected as an A-to-G change. RNA editing has been reported to exert multifaceted effects across cancers.[3-7] Site-specific recoding can alter protein function (e.g., *AZIN1*); editing can reshape splicing and miRNA biogenesis/targeting; and pervasive editing of endogenous double-stranded RNAs modulates innate immune sensing. Consequently, ADAR activity can be either oncogenic or tumor-suppressive and can influence responses to immunotherapies and epigenetic agents. While RNA editing has been implicated across many cancers—including AML—its roles in AML remain incompletely defined.

In AML specifically, evidence to date is limited. An early work showed that RNA hyperediting at a pre-mRNA branch point in *PTPN6* (SHP-1) disrupts splicing, yielding an intron-retaining transcript that is more abundant at diagnosis than in remission.[8] A recent bulk RNA-seq analysis of large AML cohorts cataloged thousands of editing events, demonstrated that higher global (Alu) editing predicts worse survival, and showed that editing burden varies across genetic subtypes.[9] Another mechanistic study revealed that core-binding factor (CBF) fusions repress *ADARB1* transcription, that *ADARB1* acts as an editing-dependent tumor suppressor in CBF-AML, and that restoring *ADARB1* activity or edited target isoforms constrains leukemogenic growth.[10] Together, these studies establish RNA editing as a contributor to AML pathobiology but leave critical questions unresolved regarding cell-type specificity and post-treatment dynamics, which are largely obscured by bulk transcriptomic analyses. In particular, it remains unknown whether RNA editing exhibits clinically informative, cell type–resolved patterns across disease states that are not captured by bulk measurements.

To address these gaps, we analyzed the single-cell RNA-seq dataset of van Galen et al.,[11] which delineates malignant and healthy hematopoietic states and includes longitudinal sampling across clinical stages. The cohort comprises well-annotated healthy and leukemia-like cell identities spanning healthy donors (HL), newly diagnosed AML (ND), and post-treatment time points—remission (RM) and persistent disease with >15% blasts (PO)—providing an ideal foundation to disentangle condition- and lineage-dependent editing programs.[11]

Here, we integrate sensitive RNA editing discovery with cell type–resolved analyses and advanced statistical modeling to chart the AML editing landscape across conditions (HL, ND, RM, PO) and lineages. Specifically, we construct a stringent catalog of recurrent edits at the pseudo-bulk level; (2) resolve cell type- and condition-specific editing changes; (3) relate global editing to *ADAR1/ADARB1* expression and nominate additional candidate regulators; and (4) assess the potential of editing features, particularly site-specific changes within key lineages, as biomarkers for disease state and prognosis, while simultaneously revealing RNA editing as an additional, cell type–specific regulatory layer underlying AML biology.

## Methods

### Cohort and study design

We analyzed 42 human scRNA-seq samples from 16 patients with AML and 5 healthy donors (GEO BioProject: PRJNA477870). Samples were grouped as healthy (HL), newly diagnosed AML (ND), and post-treatment. Post-treatment samples with ≥15% blasts were assigned to PO; the remaining post-treatment samples were assigned to RM. After quality control (see below), de-duplication (removed one cell line sample and one duplicate sample), and a <50 editing sites filter, 27 samples from 24 individuals remained: HL (n=5), ND (n=10), RM (n=7), PO (n=3) (Supplemental Table 1).

### Cell processing and RNA editing sites discovery at pseudo-bulk and cell-type level

Mononuclear cells from bone marrow (BM) aspirates of normal donors and AML patients, along with primitive cells from a fresh BM aspirate (BM5), were obtained and subjected to full-length cDNA sequencing using a high-throughput nanowell-based protocol (Seq-Well), as previously described by van Galen *et al*.[11] Owing to the technical challenges inherent to RNA editing site discovery,[12-14] we developed a comprehensive computational pipeline incorporating dual aligners with dual references, multiple stringent filtering steps, and recurrence-based site selection. RNA editing sites were first identified at the pseudo-bulk level and subsequently quantified at single-cell and cell-type–resolved levels. Full methodological details are provided in the Supplementary Methods.

### Differential RNA editing and gene expression analyses

Because RNA editing measurements derived from sequencing data are inherently sparse, overdispersed, and subject to repeated sampling within individuals, we employed a regression-based read-count modeling framework specifically designed to address these features for differential RNA editing analysis. For each editing site, edited and total read counts were modeled as a function of clinical condition using beta-binomial regression, with subject-specific random effects included when applicable. Pairwise differences in editing proportions between conditions were estimated using model-based contrasts with delta-method inference. Statistical significance was defined using false discovery rate (FDR at 0.1) together with a minimum effect-size threshold (0.05). The same modeling strategy was applied independently within each cell type to identify lineage-specific differential editing sites.

Pseudo-bulk gene expression profiles were generated by aggregating RNA-seq or scRNA-seq read counts at the sample and cell-type levels. Differential expression was assessed using quasi-likelihood negative binomial regression, with normalization and filtering performed using edgeR.[15] Models included clinical condition and, where estimable, individual-level effects to control for repeated measurements. Gene expression contrasts followed the same clinical group comparisons used in RNA editing analyses.

Full model formulations, filtering criteria, and implementation details were provided in the Supplementary Methods.

### Association between global RNA editing and gene expression

Global RNA editing activity was quantified at the pseudo-bulk level using a weighted editing proportion, defined per sample as the ratio of total edited reads to total reads summed across all high-confidence RNA editing sites (Supplementary Methods). Associations between global RNA editing and gene expression were evaluated using both correlation and regression-based frameworks.

First, Pearson correlations were computed between the weighted global editing proportion and expression of *ADAR* and *ADARB1*, with multiple testing controlled using the Benjamini–Hochberg (BH) procedure (FDR at 0.1). Second, to assess gene-wise associations while accounting for count structure and disease condition, global editing counts were modeled using beta-binomial regression with gene expression as a predictor and clinical condition as a covariate. Interaction models were further used to test for condition-dependent effects and to derive condition-specific associations. Genes with BH-adjusted FDR < 0.1 were considered statistically significant.

Full model specifications and inference procedures are described in the Supplementary Methods.

## Results

### Recurrent pseudo-bulk RNA editing in AML highlights immune/inflammatory and stress-response programs

Using a stringent discovery pipeline, we identified 2,875 RNA editing sites recurrent in ≥2 of 42 samples derived from 16 AML patients and 5 healthy donors. All identified sites were A-to-I RNA editing events, with no high-confidence C-to-U sites detected, and all overlapped entries in REDIportal v3.0, a well-established RNA editing database.[16] Of these, 697 also overlapped the AML editing sites reported by Meduri *et al*.;[9] thus, 2,178 sites were not reported in that study (Supplemental Table 2). The 2,875 sites mapped to 1,033 genes and were predominantly located in introns and 3’ UTRs (Fig. 1A), consistent with prior reports.[9, 17, 18] Gene Ontology biological process (GO:BP) enrichment of all edited genes revealed a broad functional landscape dominated by immune and inflammatory regulation, including interferon signaling, NF-κB pathways, and leukocyte activation, in line with core AML biology. In addition to immune programs, enriched terms reflected widespread regulatory functions, encompassing intracellular signal transduction (e.g., MAPK, PI3K–AKT, Rho, and Wnt pathways), transcriptional and RNA-processing regulation (including miRNA pathways), proteostasis (ubiquitination, SUMOylation, and autophagy), membrane trafficking and cytoskeletal remodeling, as well as tight control of the cell cycle, DNA damage responses, and apoptosis (Fig. 1B and Supplementary Table 3). Together, these results indicate that recurrent RNA editing sites preferentially map to genes involved in immune signaling, RNA metabolism, and cellular stress–related processes.

**Figure 1.**
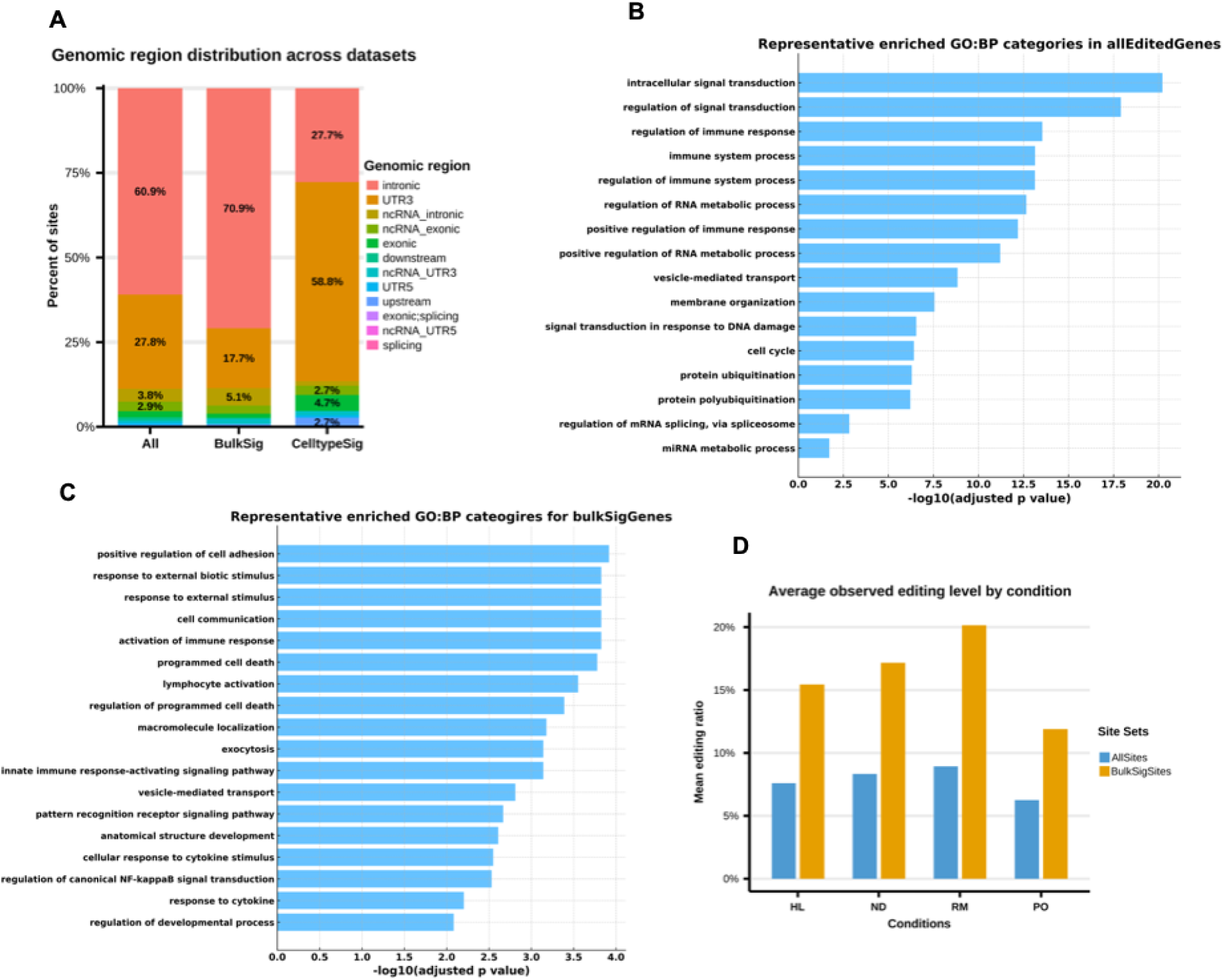
Distribution of RNA-editing sites and enriched functions. (A) Genomic locations of all discovered sites (All), differentially edited sites from the pseudo-bulk analysis (BulkSig), and cell-type–specific differentially edited sites (CelltypeSig). (B) Representative enriched functions for genes harboring all editing sites. (C) Representative enriched functions for genes with pseudo-bulk differentially edited sites. (D) Mean RNA-editing ratios by condition, shown for all sites (blue) and for pseudo-bulk differentially edited sites (yellow).

### Condition-specific RNA editing dynamics at the pseudo-bulk level

Per-site editing proportions were modeled using a beta-binomial framework, with a binomial fallback when beta-binomial models did not converge. Pairwise contrasts among HL, ND, RM, and PO were assessed via the delta method. After requiring a minimum of five total reads in at least two samples per site, 2,874 editing sites were retained for analysis.

In total, 150 differentially edited (DE) sites were detected (Table 1 and Supplemental Table 4), mapping to 124 genes. Similar to the full catalog of edited sites, most DE sites localized to introns or 3′UTRs (Fig. 1A). In contrast to the broad regulatory landscape observed for all edited genes, DE genes showed more focused enrichment for immune and host-defense programs, including pattern-recognition receptor and Toll-like receptor signaling converging on NF-κB, regulation and activation of innate and adaptive immune responses, cytokine-driven pathways, neutrophil degranulation, and platelet/hemostasis-related processes, with additional enriched terms shown in Fig. 1C and Supplementary Table 5.

**Table 1.** The number of significantly differential edited sites.

| Contrasts | Num_Sig | Num_Up | Num_Down |
| --- | --- | --- | --- |
| ND_vs_HL | 36 | 21 | 15 |
| PO_vs_HL | 40 | 11 | 29 |
| PO_vs_ND | 69 | 11 | 58 |
| PO_vs_RM | 39 | 8 | 31 |
| RM_vs_HL | 31 | 20 | 11 |
| RM_vs_ND | 39 | 20 | 19 |

**Notes:** Up/Down refers the editing ratio is higher/lower in the first condition of each contrast. For example, in ND\_vs\_HL, 21 sites show higher editing in ND.

Across comparisons, PO vs ND yielded the largest number of differential editing sites (69; Table 1). Directionality was coherent: ND and RM generally exhibited higher editing relative to HL, whereas PO displayed lower editing than all other groups, indicating that PO has the lowest global editing among the four conditions (Table 1). Mean editing proportions corroborated these patterns: RM showed the highest mean editing ratio across all the sites and all the significant subset, whereas PO showed the lowest (Fig. 1D). Notably, the mean editing ratio across significant sites exceeded the mean ratio across all sites, suggesting a greater potential functional impact of DE sites than all the sites (Fig. 1D).

Functional enrichment was broadly consistent across contrasts (e.g., ND vs HL, PO vs HL, PO vs ND, PO vs RM, RM vs HL), frequently implicating responses to external stimuli and cell adhesion. Some contrasts showed additional specificity; for example, PO vs ND and PO vs RM were enriched for vesicle-mediated transport, indicating context-dependent functional patterns (Supplemental Tables 6-11).

### Correlation between *ADAR/ADARB1* expression and global RNA editing at the pseudo-bulk level

We profiled expression of the three ADAR family members (*ADAR, ADARB1, ADARB2*) using three pseudo-bulk strategies (Methods): (1) RSEM-based quantification on aggregated reads per sample; (2) aggregation of single-cell raw counts; and (3) the mean of SCT-transformed single-cell expression. *ADARB2* was not detected and was excluded from further analyses. *ADAR* and *ADARB1* expression were highly concordant across methods (Supplemental Fig. 1); we therefore used Fragments Per Kilobase of transcript per Million mapped fragments (FPKM) values from RSEM for downstream analyses.

Pseudo-bulk expression (FPKM) distributions were broadly consistent across conditions (Fig. 2A and Supplemental Table 12). Neither *ADAR* nor *ADARB1* was differentially expressed among HL, ND, and RM; however, *ADAR* expression was significantly elevated in PO relative to other groups, whereas *ADARB1* was not. Prior studies reported no significant *ADAR* difference between HL and ND but higher *ADARB1* in healthy individuals,[9] in agreement with our findings.

**Figure 2.**
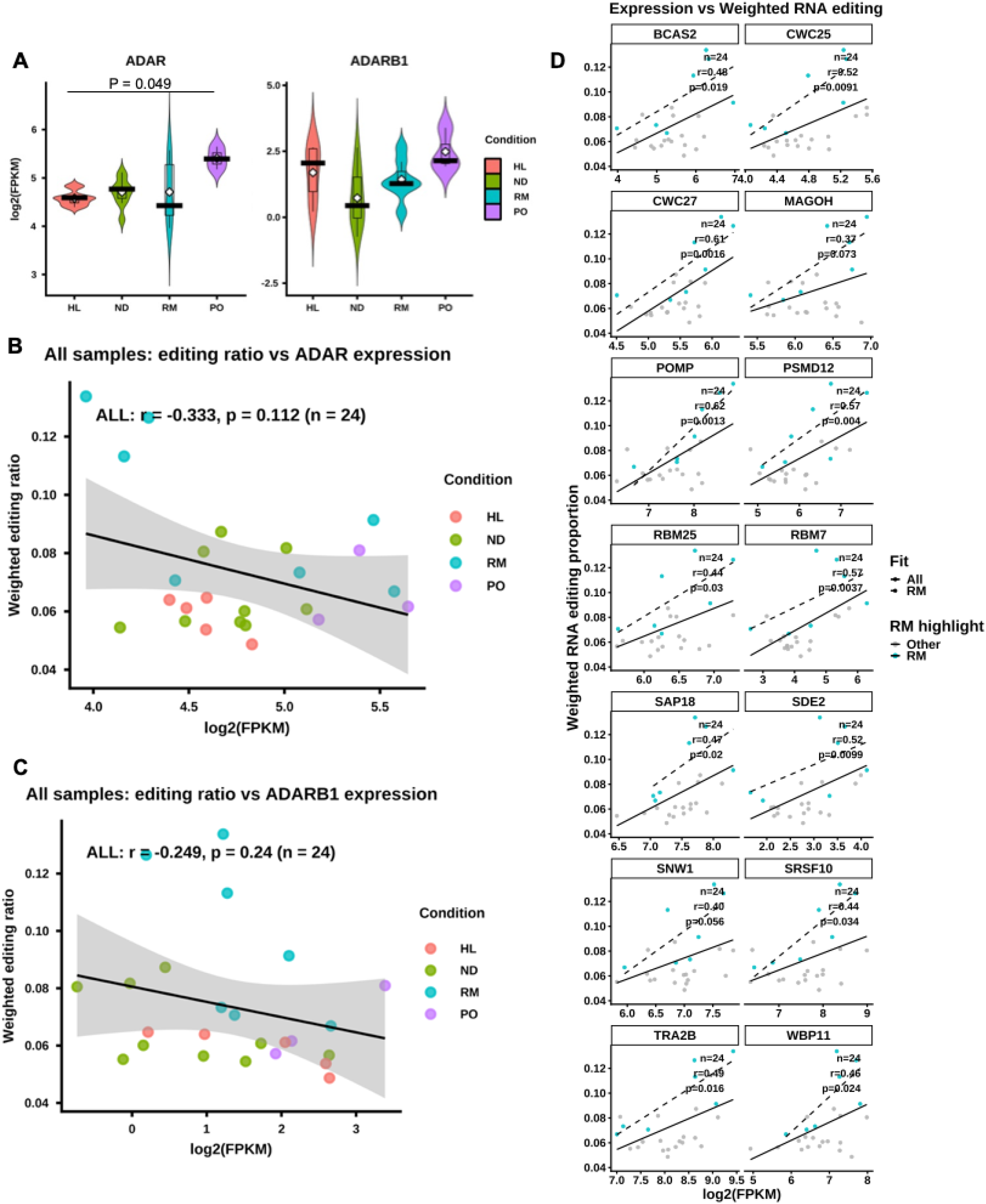
Expression of ADAR and ADARB1 and their correlation with global RNA-editing activity. (A) Expression levels of *ADAR* (left) and *ADARB1* (right) across the four clinical conditions. *ADAR* expression differed significantly between PO and HL, whereas no significant differences were observed for *ADARB1*. (B–C) Correlations between gene expression and the weighted global RNA-editing ratio, calculated from all detected sites in the pseudo-bulk analysis. (B) *ADAR* vs. weighted global RNA-editing ratio; (C) *ADARB1* vs. the same ratio. (D) Representative examples showing relationships between gene expression and the weighted global RNA-editing ratio.

Global editing per sample was defined as the fraction of edited reads across all sites (edited reads / total reads; see Methods and Supplementary Methods). At the pseudo-bulk level, *ADAR* and *ADARB1* expression both demonstrated a negative correlation with global editing; similar trends were observed when restricting to the set of significant differential edited sites (Fig. 2B-2C; Supplemental Fig. 2A-2B). This inverse association between *ADAR*/*ADARB1* expression and global editing was further confirmed independently using regression-based modeling (P = 0.011 for *ADAR* and P = 0.018 for *ADARB1*; see next section). Stratified analyses by condition largely retained the negative tendency with exceptions: in PO, both *ADAR* and *ADARB1* showed positive correlations with either all the editing sites or significantly differential edited sites (Supplemental Fig. 2C-2F)). Furthermore, Within ND, *ADAR*—but not *ADARB1*—exhibited a slight positive correlation with editing levels, consistent with prior observations (Supplemental Fig. 2C-2F).[9]

### Genes associated with global RNA editing

Given the overall negative association between global RNA editing levels and ADAR/ADARB1, we sought to identify additional candidate regulators by performing gene-wise association analyses between the per-sample global RNA editing proportion and pseudo-bulk gene expression using three beta-binomial regression models (see Methods and Supplementary Methods).

Consistent with correlation analyses, both *ADAR* and *ADARB1* exhibited significant negative associations with global RNA editing independent of disease condition, confirming the robustness of these findings (Supplemental Table 13). At FDR ≤ 0.10, we identified 2,613 genes significantly associated with global RNA editing, including 555 positively and 2,058 negatively associated genes. In addition, 282 genes demonstrated significant gene expression–by–condition interactions, indicating disease state–dependent modulation of editing–expression relationships.

Condition-specific analyses revealed pronounced differences across disease states (Supplemental Table 14). In ND, 69 genes were significantly associated with global RNA editing (20 positive, 49 negative). In contrast, RM samples exhibited a substantially stronger signal, with 515 positively and 1,973 negatively associated genes, including both *ADAR* and *ADARB1*, whereas no significant associations were detected in the remaining conditions. Thus, the global editing–associated gene landscape is largely driven by the remission condition and is dominated by negative associations.

Functional enrichment analysis highlighted distinct biological themes. The 20 positively associated genes in ND were enriched for the proteasome pathway, driven by *PSMD12* and *POMP*, which are involved in proteasome assembly and protein degradation (Supplemental Table 15). In RM, the 515 positively associated genes were strongly enriched for RNA processing pathways, particularly mRNA splicing, followed by RNA and protein metabolism, cellular stress responses, deubiquitination, PERK-regulated gene expression, and mitochondrial respiratory electron transport (Supplemental Table 16). Together, these results indicate that global RNA editing is tightly coupled to spliceosome activity and RNA metabolism, while intersecting with proteostasis, stress signaling, and cellular energy homeostasis.

Despite the marked differences in the number of associated genes between ND and RM, all positively and negatively associated genes identified in ND were also present among the corresponding RM gene sets, demonstrating strong directional consistency across disease stages. Representative correlations for genes from the proteasome and splicing pathways are shown in Fig. 2D. These associations suggest that global RNA editing reflects coordinated transcriptional programs linked to treatment response rather than static disease state.

### Cell type– and condition–specificity of global A-to-I RNA editing

We quantified RNA editing at the cell-type level using scRNA-seq-defined identities from Meduri *et al*.*[11]* Fifteen canonical hematopoietic/immune cell types were resolved, including Hematopoietic stem cell (HSC), Progenitor (Prog), Granulocyte-macrophage progenitor (GMP), Promonocyte (ProMono), Monocyte (Mono), Conventional dendritic cell (cDC), Plasmacytoid dendritic cell (pDC), Early erythroid progenitor (earlyEry), Late erythroid progenitor (lateEry), Progenitor B cell (ProB), Mature B cell (B), Plasma cell (Plasma), Naïve T cell (T), Cytotoxic T Lymphocyte (CTL), Nature Killer cell (NK). In addition, six cancer-like (leukemia-associated) states were identified: cDC-like, GMP-like, HSC-like, Mono-like, Prog-like, and ProMono-like.

Cell-type composition differed markedly by condition (Fig. 3A). HL was enriched for HSCs and progenitors; ND for leukemia-associated states (Mono-like, cDC-like, Prog-like, GMP-like, ProMono-like); RM for Mono, NK, and T cells; and PO for T, Prog-like, and NK cells.

**Figure 3.**
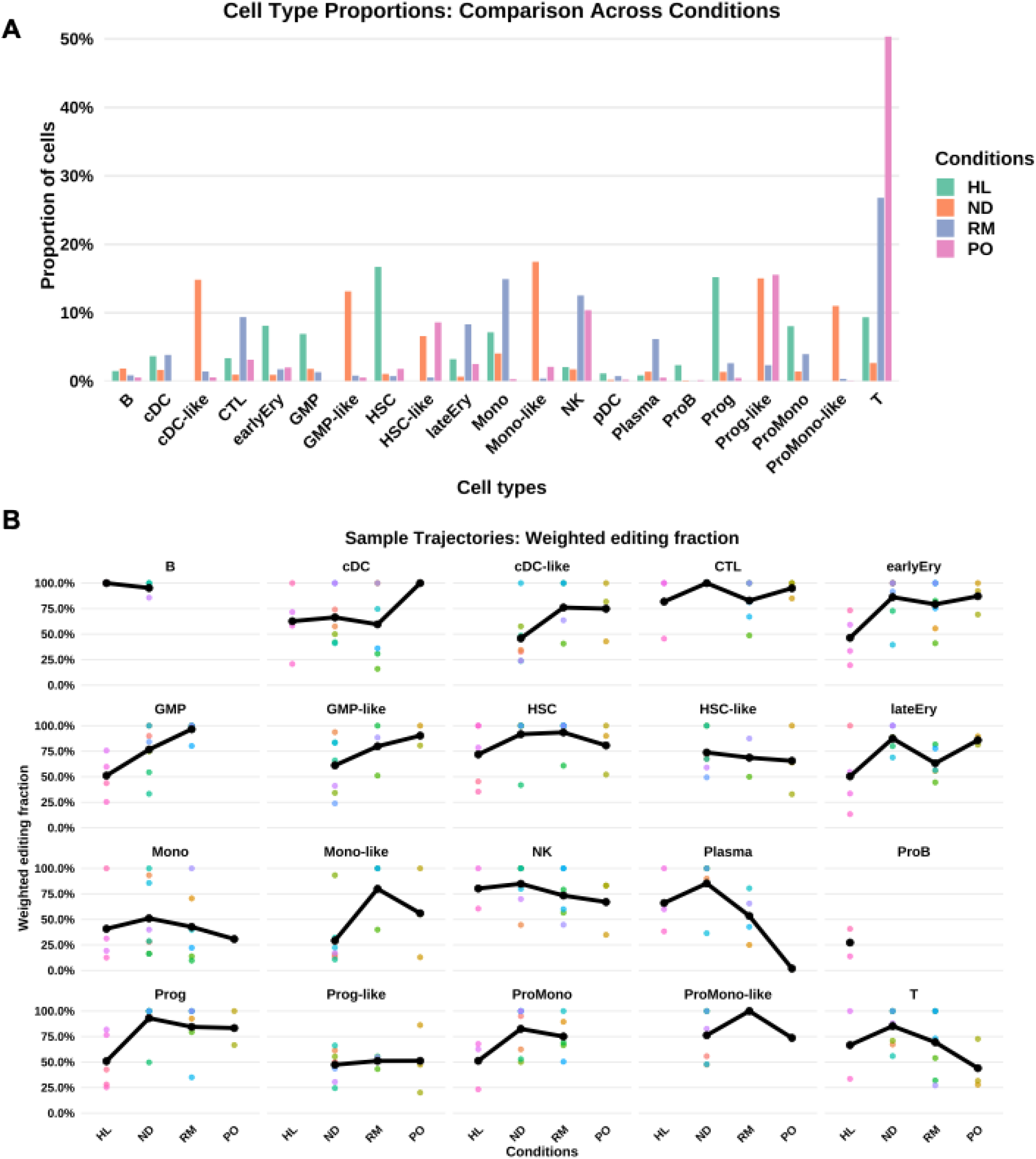
Cell-type distribution and RNA-editing dynamics across clinical conditions. (A) Cell-type composition by condition; bars indicate the proportion of each cell type within each condition. (B) Trajectories of weighted global RNA-editing levels across conditions within each cell type.

Editing-site density (number of RNA editing sites per 100 cells) also varied by cell type and condition. LateEry showed the highest mean density in HL and RM; ND showed less variability across cell type with the highest mean in cDC-like; and in PO, ProMono-like contained the highest mean editing sites (Supplemental Fig. 3A).

We next assessed the weighted editing proportion, defined as the ratio of edited to total reads aggregated across all sites within each cell type and condition. In HL, B, CTL, NK and HSC had the highest ratios. Notably, in ND, all the “cancer-like” states exhibited lower editing ratios than their healthy counterparts (e.g., cDC > cDC-like; GMP > GMP-like; HSC > HSC-like; Mono > Mono-like; Prog > Prog-like; proMono > proMono-like), indicating consistently reduced editing in leukemia-like lineages at diagnosis (Supplemental Fig. 3B). In contrast, in RM and PO, Mono-like cells displayed higher editing ratios than Mono cells.

Across conditions, ND showed the highest ratios in lateEry, Plasma, Prog, and T, and the lowest in cDC-like, GMP-like, and Mono-like. RM had the highest ratios in GMP, Mono-like, and ProMono-like. HL tended to have the lowest ratios in earlyEry, GMP, lateEry, Prog, and ProMono; ProB editing was observed only in HL, and B cells showed the highest editing ratio in HL. PO showed the highest ratios in cDC and GMP-like, and the lowest in Plasma and T cells (Fig. 3B and Supplemental Fig. 3C).

Together, these results demonstrate cell-type specificity of RNA editing that is further modulated by clinical state, supporting RNA editing as a cell- and condition-resolved feature with potential biomarker utility in AML.

### Cell-type–specific RNA editing sites across clinical states with independent cohort validation

Differential testing of individual sites identified 148 cell-type-specific sites across conditions (Table 2), enriched for antiviral, cytokine-driven, and hematopoietic immune functions with potential regulation through MHC presentation and immune-related miRNAs (Supplemental Table 17). These sites are most abundant in 3’UTR region, followed by intronic region —a distribution that contrasts with both all detected editing sites and the pseudo-bulk DE set (Fig. 1A). Each pairwise comparison included at least one cell type with ≥10 differentially edited sites, for example, RM vs ND yielded the largest count (22 sites) in Prog-like; RM vs HL had 19 sites in Mono; and PO vs HL had 18 sites in T cells. RM vs HL featured several cell types with >10 sites (Mono, T, ProMono, lateEry), suggesting these cell types contribute most to differences between RM and HL. T cells were the only cell type with ≥5 differential sites in all comparisons, indicating that RNA editing in T cells robustly distinguishes multiple conditions. Prog-like was uniquely prominent in RM vs ND, exhibiting the largest number of differentially edited sites; smaller effects were observed in PO vs RM (8 sites) and PO vs ND (2 sites), with none in the remaining three contrasts. However, Prog-like showed no apparent difference in global editing ratio between RM and ND. These findings indicate that, in addition to cell type-specific shifts in global editing, site-specific editing changes provide additional resolution across conditions in a cell type-dependent manner.

**Table 2.** Number of significantly differentially edited sites per cell type across conditions.

| Cell_Type | ND_vs_HL | PO_vs_HL | RM_vs_HL | RM_vs_ND | PO_vs_ND | PO_vs_RM |
| --- | --- | --- | --- | --- | --- | --- |
| CTL | 1 | 2 | 2 | 0 | 0 | 0 |
| GMP | 2 | 0 | 5 | 2 | 0 | 0 |
| GMP-like | 0 | 0 | 0 | 6 | 4 | 1 |
| HSC | 6 | 3 | 6 | 2 | 2 | 0 |
| HSC-like | 0 | 0 | 0 | 0 | 1 | 1 |
| Mono | 12 | 2 | 19 | 9 | 1 | 2 |
| Mono-like | 0 | 0 | 0 | 4 | 6 | 1 |
| NK | 3 | 2 | 3 | 3 | 2 | 1 |
| Plasma | 2 | 0 | 3 | 0 | 3 | 2 |
| ProMono | 7 | 0 | 13 | 5 | 0 | 0 |
| Prog | 3 | 1 | 6 | 0 | 0 | 0 |
| Prog-like | 0 | 0 | 0 | 22 | 2 | 8 |
| T | 5 | 18 | 15 | 9 | 10 | 12 |
| cDC | 4 | 0 | 6 | 0 | 0 | 0 |
| cDC-like | 0 | 0 | 0 | 5 | 5 | 1 |
| earlyEry | 3 | 2 | 3 | 1 | 0 | 1 |
| lateEry | 6 | 2 | 11 | 3 | 2 | 0 |

**Notes:** Cells with values >15 are shaded red; values >10 are green; yellow highlights the threshold values used for coloring.

RNA-editing sites identified at the pseudo-bulk and cell-type levels overlapped significantly: 150 pseudo-bulk and 148 cell-type sites, with 19 shared (∼13% overlap; Fisher’s exact, two-sided, p ≈ 3.5×10^−^4). We highlight five sites differing significantly across conditions in pseudo-bulk and cell-type analyses (Fig. 4A-4B, Supplemental Table 18-19): three lie in 3’UTRs—*GBP4* (an interferon-inducible large GTPase in innate immunity), *SPN/CD43* (a leukocyte sialoglycoprotein regulating T-cell activation/adhesion; AML blasts often display the *CD43s* glycoform), and *TNFSF10/TRAIL* (a death-receptor ligand activating extrinsic apoptosis and inducible by HDAC inhibition)—and two are intronic—*EMB* (an Ig-superfamily adhesion molecule that promotes ECM interactions) and *FKBP5* (an Hsp90 co-chaperone modulating glucocorticoid signaling; intronic variation has been linked to cytarabine response in pediatric AML). At the cell-type level, three of these sites recapitulated the same condition contrast observed in pseudo-bulk but only within specific lineages, whereas the remaining two were significant in different contrasts. Together, these results show that while significant concordance exists between pseudo-bulk and cell-type analyses, it is largely lineage-restricted; most signals are cell-type– and condition-specific (∼87% non-overlapping), indicating bulk signals do not simply average cell-type effects.

**Figure 4.**
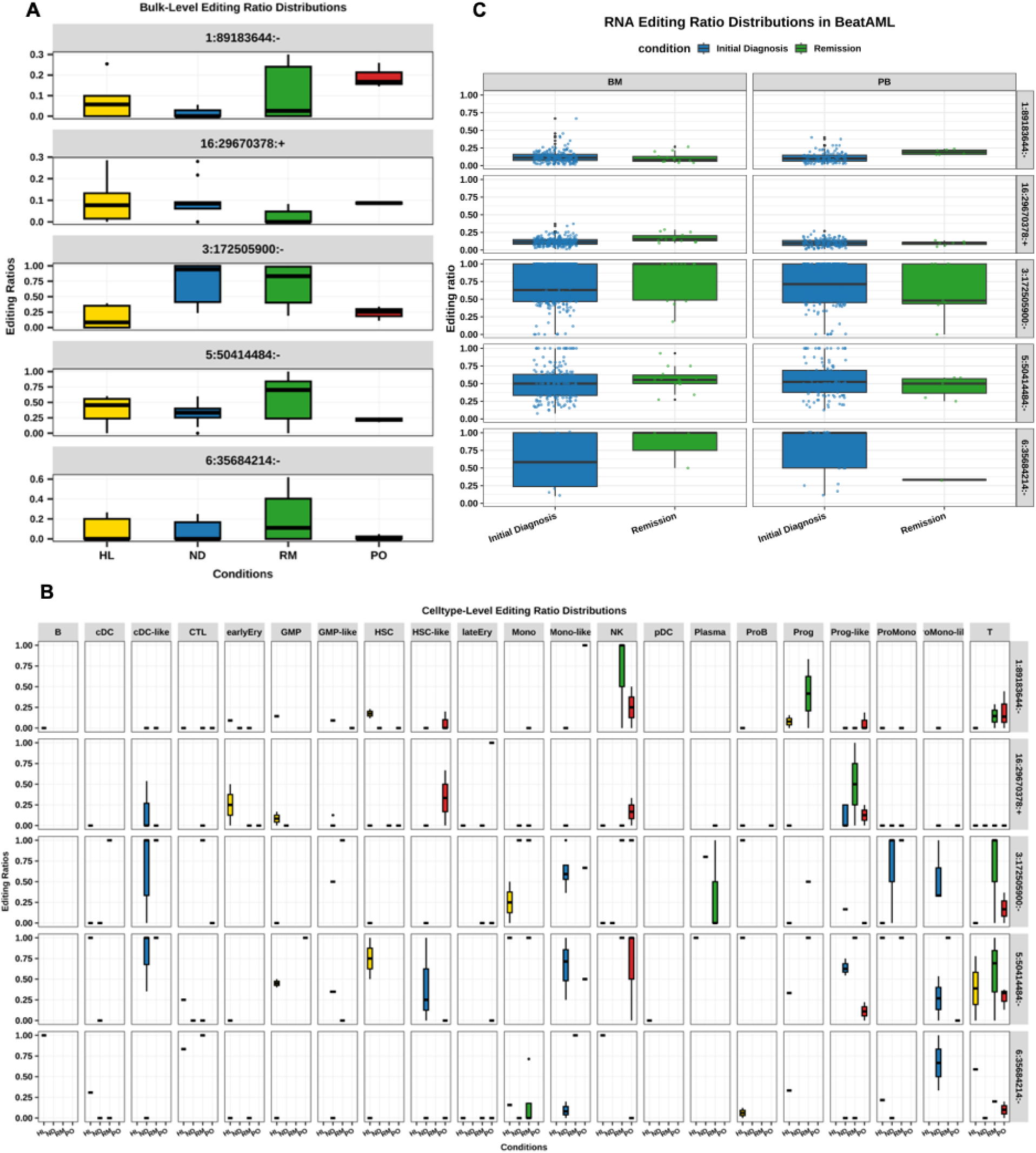
Representative RNA-editing sites across clinical conditions. (A) Pseudo-bulk editing ratios for five loci across HL, ND, RM, and PO: chr1:89 183 644 (–), 3’UTR of *GBP4*; chr16:29 670 378 (+), 3’UTR of *SPN*; chr3:172 505 900 (–), 3’UTR of *TNFSF10*; chr5:50 414 484 (–), intronic *EMB*; chr6:35 684 214 (–), intronic *FKBP5*. Comparisons showing significant differences for these five sites are provided in Supplementary Table 18. (B) Cell-type–resolved editing ratios for the same five sites across the four conditions. Detailed comparison results are provided in Supplementary Table 19. (C) Distribution of editing ratios for the same five sites in the BEATAML cohort across two clinical conditions in both bone marrow and peripheral blood samples.

We next evaluated whether these five candidate RNA editing sites could be independently validated in two external cohorts, The Cancer Genome Atlas Acute Myeloid Leukemia (TCGA-LAML) and the Beat Acute Myeloid Leukemia Master Trial (BEATAML) (Supplementary Methods). All five sites were detectable in both datasets. Because BEATAML includes clinical states analogous to those examined in this study—initial diagnosis (corresponding to ND) and remission (corresponding to RM)—we focused subsequent validation analyses on this cohort.

BEATAML comprises two specimen types, bone marrow aspirates (BM; n = 296) and peripheral blood (PB; n = 246), which were analyzed separately. At both the pseudo-bulk and cell-type–resolved levels, three of the five sites—16:29670378:+ (3′UTR of *SPN*), 5:50414484:− (intronic region of *EMB*), and 6:35684214:− (intronic region of *FKBP5*)—showed significant differential editing between ND and RM (Supplemental Table 18-19). Notably, two sites (5:50414484:− and 6:35684214:−) exhibited consistent directionality between initial diagnosis and remission in BM samples, matching the tissue context of our single-cell dataset (Fig. 4C). Notably, we observed reversed directionality for all the five sites in peripheral blood, suggesting tissue-specific regulation of RNA editing.

The third site, 16:29670378:+, displayed higher editing in ND at the pseudo-bulk level but higher editing in remission in BEATAML overall; however, this pattern was concordant with the cell-type–specific results observed in progenitor-like cells (Supplemental Table 19 and Fig. 4B). In addition, a fourth editing site, 3:172505900:− located in the 3′UTR of *TNFSF10*, demonstrated significantly higher editing levels in RM compared with ND in the cell-type–resolved analysis (Supplemental Table 19 and Fig. 4B). The same directional change was observed in BEATAML BM samples (Fig. 4C), further underscoring the robustness and lineage dependence of this signal.

Finally, we assessed the clinical relevance of these five editing sites by testing their associations with key BEATAML clinical features, including overall survival, European LeukemiaNet (ELN) risk classification, and blast percentages in bone marrow and peripheral blood (Supplementary Methods). Each site was significantly associated with at least one clinical variable in either BM or PB samples (Supplemental Table 20), supporting the potential clinical relevance of these RNA editing events.

## Discussion

By aggregating sc RNA-seq reads to the sample and resolving edits at both the pseudo-bulk and cell-type levels, we characterize a multiscale landscape of RNA editing patterns in AML. The catalog is enriched in intronic and 3’UTR regions and maps to 1,033 genes involved in regulation of signal transduction and innate and adaptive immunity. Building on this atlas, we show that RNA editing levels differ systematically across disease states and that these differences exhibit pronounced cell-type specificity.

At the pseudo-bulk level, we identified 150 differentially edited sites across clinical states. Editing directionality was coherent: most DE sites in ND and RM have higher ratios than HL, whereas PO was lower than all other groups, consistent with its lowest mean global editing. In contrast, RM displayed the highest mean editing ratio (both across all sites and the DE subset). Notably, DE sites consistently exhibited higher mean editing ratios than the full set of sites across all conditions. Functional enrichment patterns were broadly shared across contrasts, while comparisons involving PO uniquely highlighted vesicle-mediated transport pathways. Collectively, these findings indicate condition-specific differences in RNA editing burden, with persistent disease showing distinct enrichment patterns that implicate vesicle-mediated transport–related pathways.

Dissecting by cell identity revealed pronounced heterogeneity. Several leukemia-associated states exhibited lower global editing than their healthy counterparts in ND, indicating disease-linked suppression of editing within specific lineages in the newly diagnosed AML. Site-level testing identified 148 cell type–specific DE sites, enriched for immune response, response to external stimuli, and Golgi calcium ion transport, pointing to editing changes at transcripts that could modulate antigen presentation, signaling, and secretory stress responses.

Three lines of evidence in our dataset highlight the value of cell-type–resolved analysis. (1) T cells harbored five or more DE sites in every clinical contrast, positioning T-cell editing as a sensitive discriminator of disease state. Prog-like cells contributed the largest DE-site burden in the RM vs ND comparison despite no detectable difference in global editing, indicating that cell-restricted, site-specific changes—not global shifts—encode condition information. (3) DE-site overlap between cell-type and pseudo-bulk was small even though significant; when sites were shared, their behavior remained cell-type and condition specific. Together, these findings demonstrate that bulk measures obscure informative lineage-specific editing programs and that cell-type granularity provides additional resolution for biological interpretation and biomarker prioritization.

Across three pseudo-bulk quantification strategies, *ADAR* and *ADARB1* expression estimates were concordant. Unexpectedly, both showed overall negative associations with global editing. Prior studies reported positive associations in ND, yet *ADAR* was not differentially expressed between ND and HL despite higher global editing, and *ADARB1* was up-regulated in HL without correlating with editing levels. These observations indicate that *ADAR/ADARB1* transcript abundance alone does not explain global editing variation in AML. Possible explanations include isoform- or localization-specific regulation, residual compositional effects after aggregation—supported by condition-stratified trends—and regulation by additional editing cofactors.

Comparisons between HL and RM showed marked differences in both RNA editing and gene expression (Table 1; Supplemental Table 21; Supplemental Fig. 4A-4B), indicating that although having achieved remission after treatment, RM has not fully reverted to the HL state. Hierarchical clustering placed RM closest to HL among disease groups by expression (Supplemental Fig. 4A), whereas RNA-editing profiles showed a mixed proximity pattern (Supplemental Fig. 4B), suggesting that these modalities capture overlapping but distinct aspects of post-treatment biology. Together, these results support the use of both RNA editing and gene expression as complementary biomarkers for clinical monitoring.

A limitation of this study is the reliance on a single published single-cell RNA-seq dataset, reflecting the current scarcity of datasets that jointly provide sufficient read depth, cell-type resolution, and longitudinal clinical annotation for robust RNA editing analysis. To mitigate this limitation, we applied multiple complementary and orthogonal analytical strategies—including pseudo-bulk aggregation, cell-type–resolved modeling, and site-specific regression analyses—that consistently converged on the same biological conclusions. The concordance of results across these analytical scales provides strong internal validation of the observed RNA editing patterns, despite the use of a single single-cell cohort. As additional single-cell datasets with adequate sequencing depth and clinical annotation become available, future studies will be well positioned to further validate and extend these findings.

In addition, we did not resolve ADAR isoforms or post-translational regulatory mechanisms, which may critically influence catalytic activity and editing specificity. The lack of matched DNA sequencing data may also lead to misclassification of rare genomic variants as RNA editing events or, conversely, exclusion of true edits under stringent single-nucleotide polymorphism filtering criteria.

## Conclusions

In summary, this study provides a condition- and cell-type–resolved map of RNA editing in AML, reveals overall negative relationships between global editing and *ADAR/ADARB1* expression, and nominates site-specific, lineage-restricted editing events—particularly in T cells and progenitor-like cells—as promising mechanistic insights and translational targets for future investigation.

## List of abbreviations

AML: Acute myeloid leukemia
BEATAML: Beat Acute Myeloid Leukemia Master Trial
BM: Bone marrow
B: Mature B cell
cDC: Conventional dendritic cell
CTL: Cytotoxic T lymphocyte
DE: Differentially edited
ELN: European LeukemiaNet
earlyEry: Early erythroid progenitor
FDR: False discovery rate
FPKM: Fragments per kilobase of transcript per million mapped fragments
GMP: Granulocyte–macrophage progenitor
HL: Healthy donors
HSC: Hematopoietic stem cell
lateEry: Late erythroid progenitor
Mono: Monocyte
ND: Newly diagnosed acute myeloid leukemia
NK: Natural killer cell
pDC: Plasmacytoid dendritic cell
Plasma: Plasma cell
PM: Peripheral blood
PO: Persistent disease
Prog: Progenitor
ProB: Progenitor B cell
ProMono: Promonocyte
RM: Remission
scRNA-seq: single cell RNA sequencing
T: Naïve T cell
TCGA-LAML: The Cancer Genome Atlas Acute Myeloid Leukemia

## Declarations

### Ethics approval and consent to participate

All data was de-identified and derived from publicly available datasets. No new patient data was presented in this body of work.

### Consent for publication

Not applicable.

### Availability of data and materials

The scRNA-seq data and associated clinical annotations analyzed in this study are publicly available through the Gene Expression Omnibus (GEO) under BioProject accession PRJNA477870. Bulk RNA-seq data from independent validation cohorts are available through dbGaP for TCGA-LAML (phs000178) and BEATAML (phs001657), subject to controlled-access approval. All data processing steps, analysis workflows, and parameters are described in the Methods and Supplementary Methods.

### Authors’ contributions

T.G. conceived the study and performed analysis. D.B. refined the RNA editing discovery pipeline. T.G. drafted the manuscript, A.E.T. and G.S.G.M. interpreted the results. All authors reviewed, edited, and approved the final version.

## Acknowledgements

The authors thank the investigators who generated and made publicly available the scRNA-seq and bulk RNA-seq datasets used in this study.

