## Supplementary Methods and Figures for "Multiscale RNA editing analysis reveals cell type–specific regulatory programs across disease states in acute myeloid leukemia"

#### 1. Stringent read preprocessing and alignment strategy

Per sample, reads from all cells were merged into one FASTQ. Read quality was assessed with FastQC v0.11.7; adapters/low-quality bases were trimmed by Trimmomatic v0.39 using "LEADING:20 TRAILING:20 SLIDINGWINDOW:4:15 MINLEN:30."[1] Reads were aligned to GRCh38 using two strategies: (1) STAR v2.7.9a (two-pass),[2] retaining uniquely mapped reads for downstream analyses; and (2) Bowtie v1.2.3[3] to both a customized splice database and GRCh38, retaining uniquely mapped reads and, for dual hits, the higher-scoring alignment. Only reads with consistent alignments between STAR and Bowtie were carried forward for RNA-editing discovery as dual-alignment strategy has been shown to reduce false-positive calls by approximately 50%.[4] For cell type–specific RNA-editing discovery and quantification, aligned reads were then partitioned by cell type.

#### 2. RNA editing site calling and filters

Candidate A-to-I sites were called from pileups with samtools (v1.18)[5] and stringently filtered to minimize technical and genomic artifacts: (1) Genetic polymorphisms: we removed sites overlapping dbSNP build 151 SNVs and all "common" human variants from NCBI (https://www.ncbi.nlm.nih.gov/variation/docs/human_variation_vcf/. (2) Problematic regions: we excluded loci within simple repeats (v10.3.28) and rRNA annotated by the UCSC RepeatMasker track. (3) Strand bias: two-sided Fisher's exact test was used to identify strand bias sites and sites with Benjamini–Hochberg (BH) FDR < 0.05 were excluded. (4) Positional bias: distance-to-read-start test and significant positional bias sites (BH FDR < 0.001) were excluded. (5) Nearby mutations: removed sites with any non-A-to-G/T-to-C mutation within ± 49 bp (Length of the reads). (6) Recurrence/coverage: retained sites observed in ≥2 samples.

#### 3. Differential RNA editing analysis at pseudo-bulk and cell-type level

**Model.** For each site *s* and sample *i*, let $y_{si}$ be edited read count, $n_{si}$ total reads, $\mu_{si}$ the mean editing ratio and $\rho_s$ the overdispersion. We fit a beta-binomial GLMM using glmmTMB:[6]

$$y_{si} \sim \text{BetaBinomial}(n_{si}, \mu_{si}, \rho_s),$$

$$\text{logit}(\mu_{si}) = \beta_0 + \beta_{\text{ND}}\, 1_{\{\text{ND}\}} + \beta_{\text{RM}}\, 1_{\{\text{RM}\}} + \beta_{\text{PO}}\, 1_{\{\text{PO}\}} + b_{\text{ind}[i]},$$

with HL as the reference level. A subject-specific intercept $b_{\text{ind[i]}} \sim \mathcal{N}(0, \sigma_b^2)$ was included when repeated measurements existed; if the beta-binomial failed to converge, we refit a binomial GLM: $y_{si} \sim \text{Binomial}(n_{si}, \mu_{si}),\ \text{logit}(\mu_{si}) = x_i^T \beta(+b_{\text{ind}[i]})$

**Contrasts and inference**. For group $L \in \{HL, ND, RM, PO\}$, the model-based proportion is $\hat{p}_L = \text{logit}^{-1}(\eta_L)$ with $\eta_L = x_L^T \hat{\beta}$. For contrast A_vs_B (order fixed as HL < ND < RM < PO), the effect is

$$\Delta_{A,B} = \hat{p}_A - \hat{p}_B.$$

The delta-method SE uses the fixed-effects covariance V: with $g = \hat{p}_A(1 - \hat{p}_A)x_A - \hat{p}_B(1 - \hat{p}_B)x_B$,

$$\mathrm{SE}(\Delta_{A,B}) \approx \sqrt{g^T V g}.$$

Two-sided P values were BH-adjusted within contrast; significance required FDR < 0.10 and $|\hat{\Delta}_{A,B}^{\mathrm{obs}}| > 0.05$ where

$$\hat{p}_L^{obs} = \frac{\sum_i y_{si}\ 1_{\{i \in L\}}}{\sum_i n_{si}\ 1_{\{i \in L\}}}, \qquad \hat{\Delta}_{A,B}^{\mathrm{obs}} = \hat{p}_A^{obs} - \hat{p}_B^{obs}.$$

We applied the same approach at the cell-type level, evaluating each site within each cell type across the four clinical conditions (HL, ND, RM, and PO).

### 4. Pseudo-bulk gene expression and differential expression analysis

We obtained per-sample pseudo-bulk counts by two approaches: 1) RNA-seq aggregation: trimmed reads were aligned with STAR v2.7.9a[2] and quantified by RSEM v1.3.3.[7] From this approach, we obtained read counts and Fragments Per Kilobase of transcript per Million mapped reads (FPKM) values for each gene for downstream analysis. 2) scRNA-seq aggregation: the GEO count matrix was summed across all cells per sample. For cell-type-specific analyses, counts were summed per cell type per sample. Samples used for pseudo-bulk required ≥20 cells.

Read counts were normalized with edgeR TMM method.[8] Counts Per Million (CPM) were obtained using the function of cpm in edgeR. Genes were filtered using filterByExpr with min.count = 10, min.total.count = 15, min.prop = 0.5. We fit quasi-likelihood negative-binomial GLMs:

$$Y_{gs} \sim \mathrm{NB}(\mu_{gs}, \phi_g), \qquad \log \mu_{gs} = \log N_s + \log f_s + X_s^{\mathrm{T}} \beta_g,$$

where $N_s$ is library size, $f_s$ the TMM normalization factor, and $X_s$ encodes conditions (and individual if estimable). Designs used were ~ 0 + conditions + indID when full-rank; otherwise ~0 + conditions. Contrasts compared ND, RM, PO to HL and the pairwise differences among ND, RM, PO. For differential expression analyses of ADAR and ADARB1, a nominal P < 0.05 was used to assess significance. For transcriptome-wide analyses, statistical significance was defined using a false discovery rate (FDR) threshold of 0.05 and a minimum of two fold change.

### 5. RNA editing and gene expression association at the pseudo-bulk level

#### 5.1. Definition of global RNA editing proportion

For each sample, global RNA editing activity was quantified using a weighted editing proportion:

$$\hat{p}_{\text{weighted}} = \frac{\sum_s y_s}{\sum_s n_s},$$

where $y_s$ and $n_s$ denote the total number of edited reads and total reads, respectively, aggregated across all high-confidence RNA editing sites $s$. This formulation preserves the contribution of individual sites while providing a single summary measure of global RNA editing activity per sample.

#### 5.2. Correlation analysis with gene expression

To assess global relationships between RNA editing activity and gene expression, we computed Pearson correlation coefficients between $\hat{p}_{\text{weighted}}$ and pseudo-bulk gene expression levels, including *ADAR*, *ADARB1*, and all expressed genes. Gene expression values were obtained from normalized RSEM FPKM estimates and $\log_2$-transformed [$\log_2$(FPKM + 0.1)]. P values were adjusted for multiple testing using the Benjamini–Hochberg (BH) procedure, and associations with BH-adjusted FDR < 0.10 were considered statistically significant.

#### 5.3. Beta-binomial regression modeling

To formally model the association between gene expression and global RNA editing while accounting for the count-based nature of editing data and disease condition, we fit gene-wise beta-binomial regression models using glmmTMB. For each sample $i$, let $y_i = \sum_s y_{si}$ denote the total number of edited reads and $n_i = \sum_s n_{si}$ the total number of reads across all sites. Editing counts were modeled as:

$$y_i \sim \text{Beta-Binomial}(n_i, p_i, \phi), \text{logit}(p_i) = \eta_i,$$

where $p_i$ represents the expected global editing proportion and $\phi$ captures overdispersion.

#### 5.4. Overall and condition-dependent associations

For each gene, we first evaluated condition-adjusted associations using nested models:

- Null model (M0):

$$\text{logit}(p_i) = \alpha + \gamma_{C_i}$$

- Additive model (M1):

$$\text{logit}(p_i) = \alpha + \beta x_i + \gamma_{C_i}$$

where $x_i$ denotes gene expression and $C_i$ indicates disease condition (HL, ND, RM, PO). Likelihood ratio tests (LRTs) comparing M1 to M0 were used to assess overall associations, with BH correction applied across genes.

To test whether gene–editing associations differed by disease condition, we fit an interaction model:

- Interaction model (M2):

$$\text{logit}(p_i) = \alpha + \beta x_i + \gamma_{C_i} + \delta_{C_i} x_i.$$

LRT comparing M2 to M1 was used to assess condition-dependent effects. For genes with significant interactions, condition-specific slopes were derived from the interaction model, and Wald tests were used to evaluate their significance. Multiple testing correction was applied separately within each condition using the BH procedure.

### 6. RNA editing sites discovery and clinical association analysis in BEATAML and TCGA-LAML

Bulk RNA-seq data were obtained from dbGaP for the TCGA-LAML (dbGaP accession number: phs000178) and BEATAML (dbGaP accession number: phs001657) cohorts. RNA editing site discovery and quantification followed the same pipeline used for pseudo-bulk analyses of the scRNA-seq dataset, including alignment, filtering, and site-level calling.

Associations between the five selected RNA editing sites and clinical features in BEATAML were evaluated using beta-binomial regression for European LeukemiaNet (ELN) risk classification and blast percentages in bone marrow and peripheral blood separately with adjusting for age, sex, and the top two principal components derived from interferon-stimulated genes (ISGs). Associations with overall survival were assessed using Cox proportional hazards regression.

#### 7. Functional enrichment analysis

Functional enrichment of edited genes used gprofiler2 with gSCS multiple-testing correction;[9] significant terms were reported at FDR < 0.05.


1. Bolger AM, Lohse M, Usadel B: **Trimmomatic: a flexible trimmer for Illumina sequence data.** *Bioinformatics* 2014, **30:**2114-2120.
2. Dobin A, Davis CA, Schlesinger F, Drenkow J, Zaleski C, Jha S, Batut P, Chaisson M, Gingeras TR: **STAR: ultrafast universal RNA-seq aligner.** *Bioinformatics* 2013, **29:**15-21.
3. Langmead B, Trapnell C, Pop M, Salzberg SL: **Ultrafast and memory-efficient alignment of short DNA sequences to the human genome.** *Genome Biol* 2009, **10:**R25.
4. Wood DE, White JR, Georgiadis A, Van Emburgh B, Parpart-Li S, Mitchell J, Anagnostou V, Niknafs N, Karchin R, Papp E, et al: **A machine learning approach for somatic mutation discovery.** *Sci Transl Med* 2018, **10**.
5. Li H, Handsaker B, Wysoker A, Fennell T, Ruan J, Homer N, Marth G, Abecasis G, Durbin R, Genome Project Data Processing S: **The Sequence Alignment/Map format and SAMtools.** *Bioinformatics* 2009, **25:**2078-2079.
6. McGillycuddy M, Popovic G, Bolker BM, Warton DI: **Parsimoniously Fitting Large Multivariate Random Effects in glmmTMB.** *Journal of Statistical Software* 2025, **112:**1 - 19.
7. Li B, Dewey CN: **RSEM: accurate transcript quantification from RNA-Seq data with or without a reference genome.** *BMC Bioinformatics* 2011, **12:**323.
8. Chen Y, Chen L, Lun ATL, Baldoni PL, Smyth GK: **edgeR v4: powerful differential analysis of sequencing data with expanded functionality and improved support for small counts and larger datasets.** *bioRxiv* 2024**:**2024.2001.2021.576131.
9. Kolberg L, Raudvere U, Kuzmin I, Vilo J, Peterson H: **gprofiler2 -- an R package for gene list functional enrichment analysis and namespace conversion toolset g:Profiler.** *F1000Res* 2020, **9**.

**Supplementary Figures**

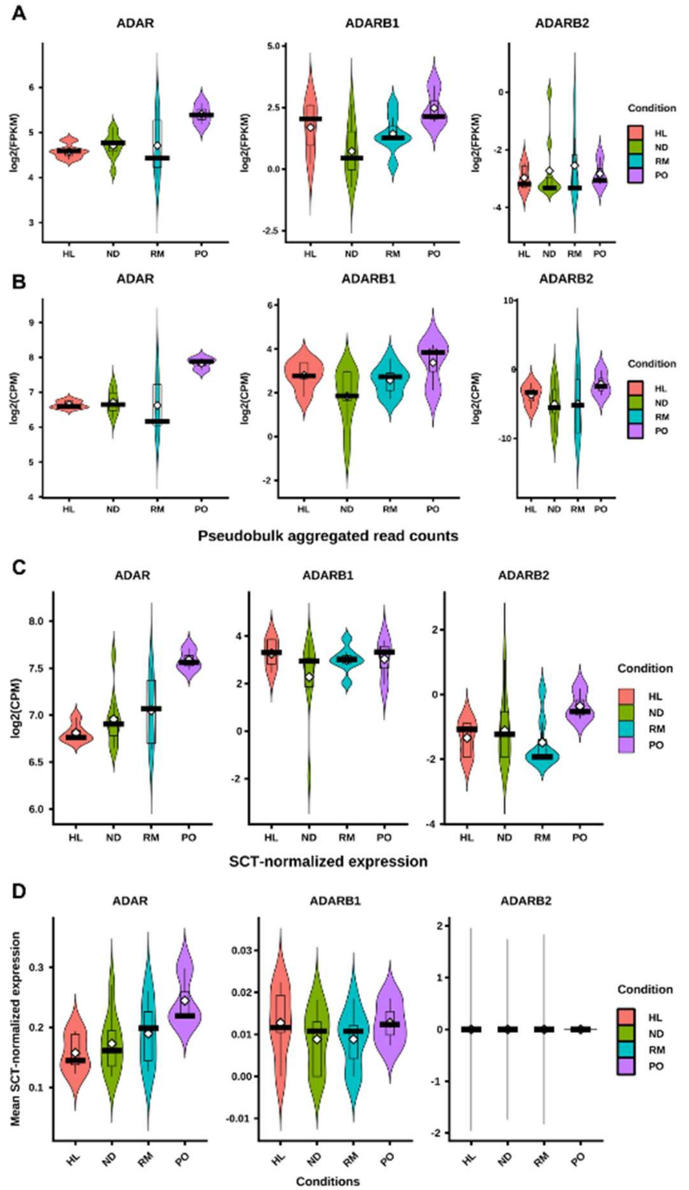


**Supplemental Figure 1. ADAR and ADARB1 expression across four clinical conditions by pseudo-bulk and single-cell quantification.** (A) RSEM-derived FPKM (pseudo-bulk of aggregated reads). (B) RSEM-derived $\log_2$(CPM) (pseudo-bulk of aggregated reads). (C) Pseudo-bulk CPM from aggregated single-cell raw counts. (D) Single-cell expression after SCT (sctransform) normalization. Abbreviations: CPM, counts per million; FPKM, fragments per kilobase of transcript per million mapped reads; SCT, sctransform normalization.

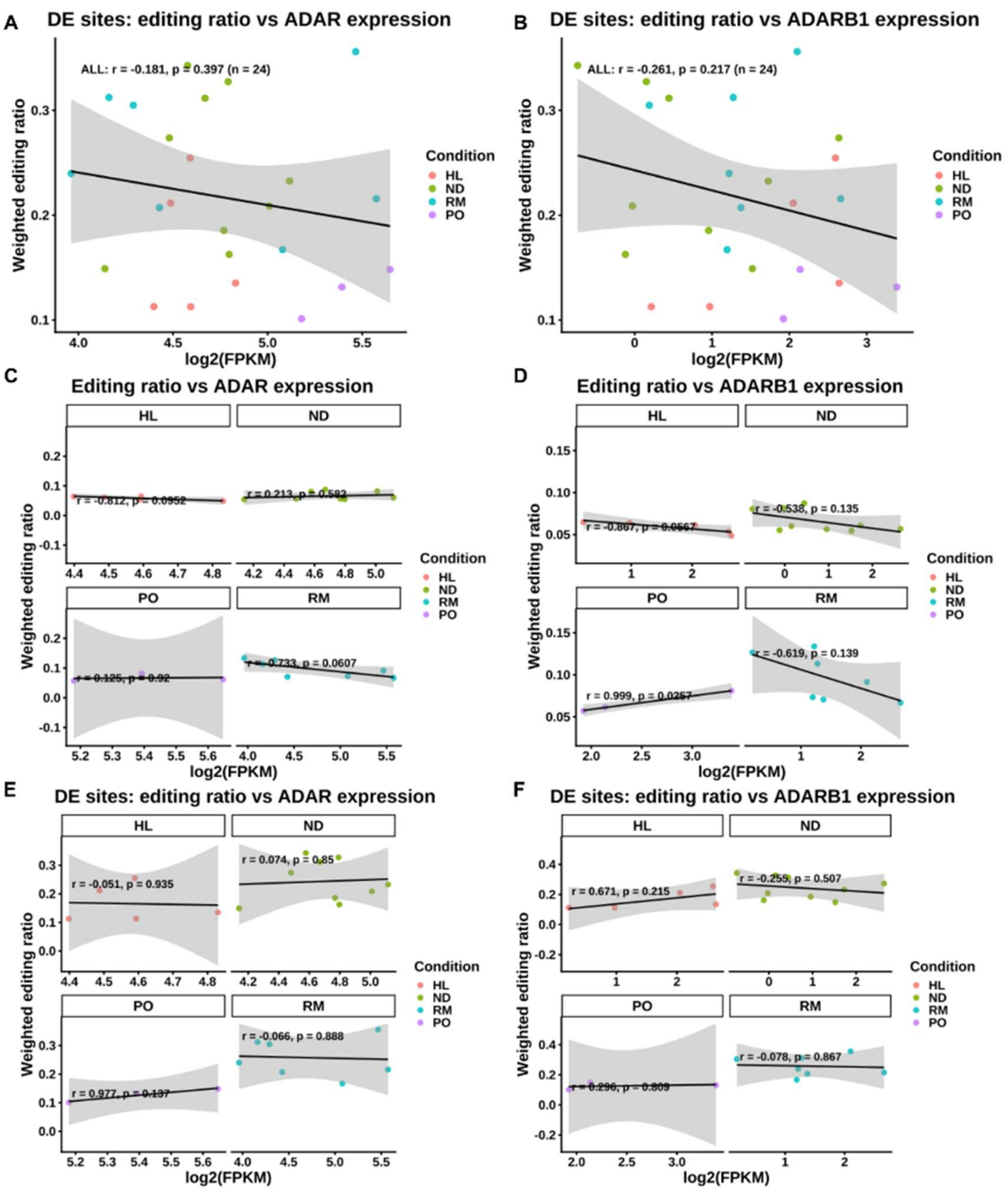


**Supplemental Figure 2. Correlation of *ADAR* and *ADARB1* expression with the weighted global RNA-editing ratio.** (A–B) Correlations between *ADAR* (A) and *ADARB1* (B) expression and the weighted RNA-editing ratio, calculated from differentially edited (DE) sites identified in the pseudo-bulk analysis. (C–D) Within-condition correlations between *ADAR* (C) and *ADARB1* (D) expression and the weighted global RNA-editing ratio, calculated from all detected sites. (E–F) Within-condition correlations between *ADAR* (E) and *ADARB1* (F) expression and the weighted RNA-editing ratio, calculated from DE sites in the pseudo-bulk analysis.

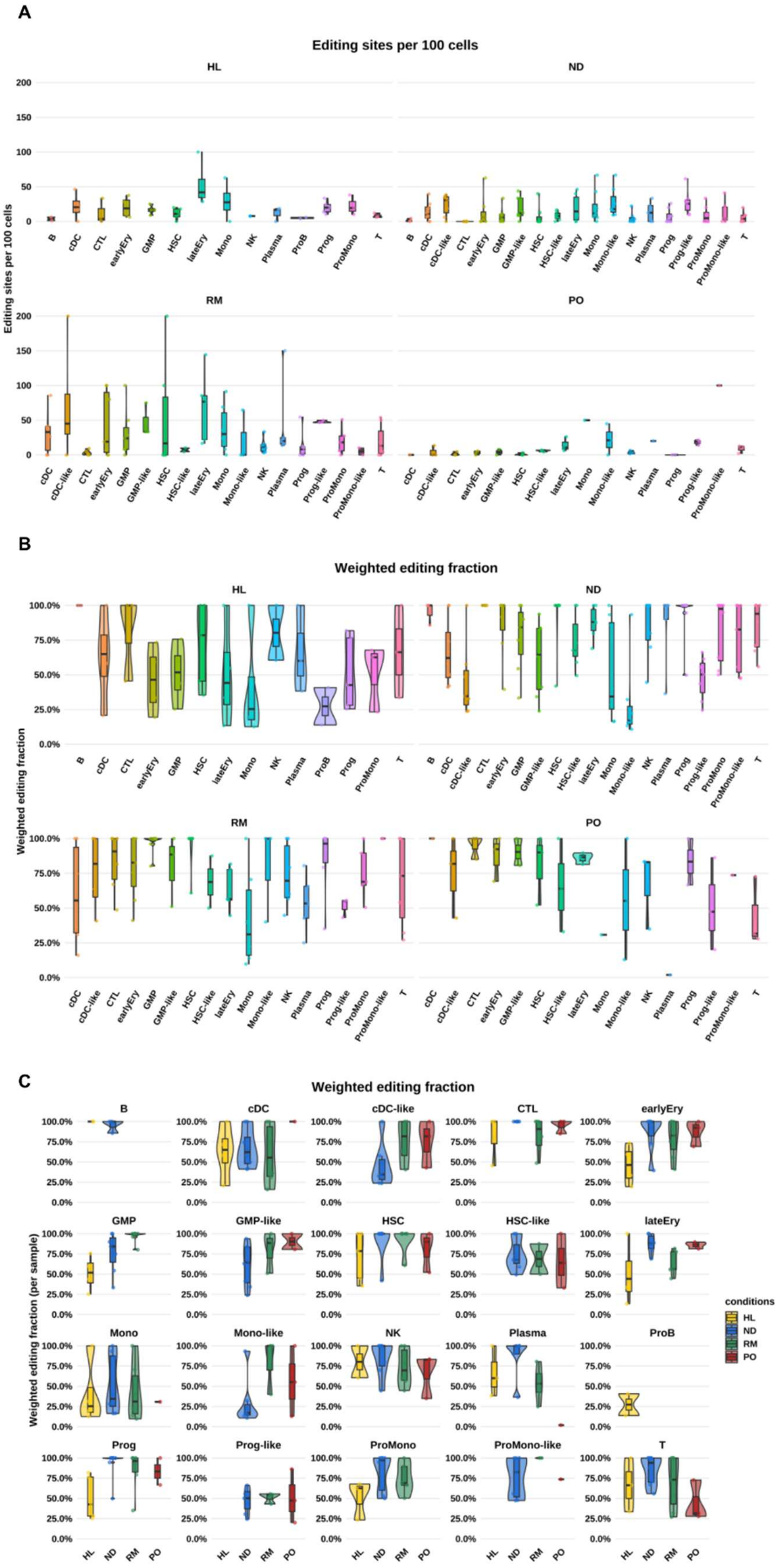


**Supplemental Figure 3. Cell-type–resolved RNA editing across clinical conditions.** (A) RNA-editing density by cell type within each condition, defined as the number of editing events per 100 cells. (B) Global RNA-editing levels by cell type within each condition (HL, ND, RM, PO). (C) Distributions of global RNA-editing levels across the four conditions within each cell type.

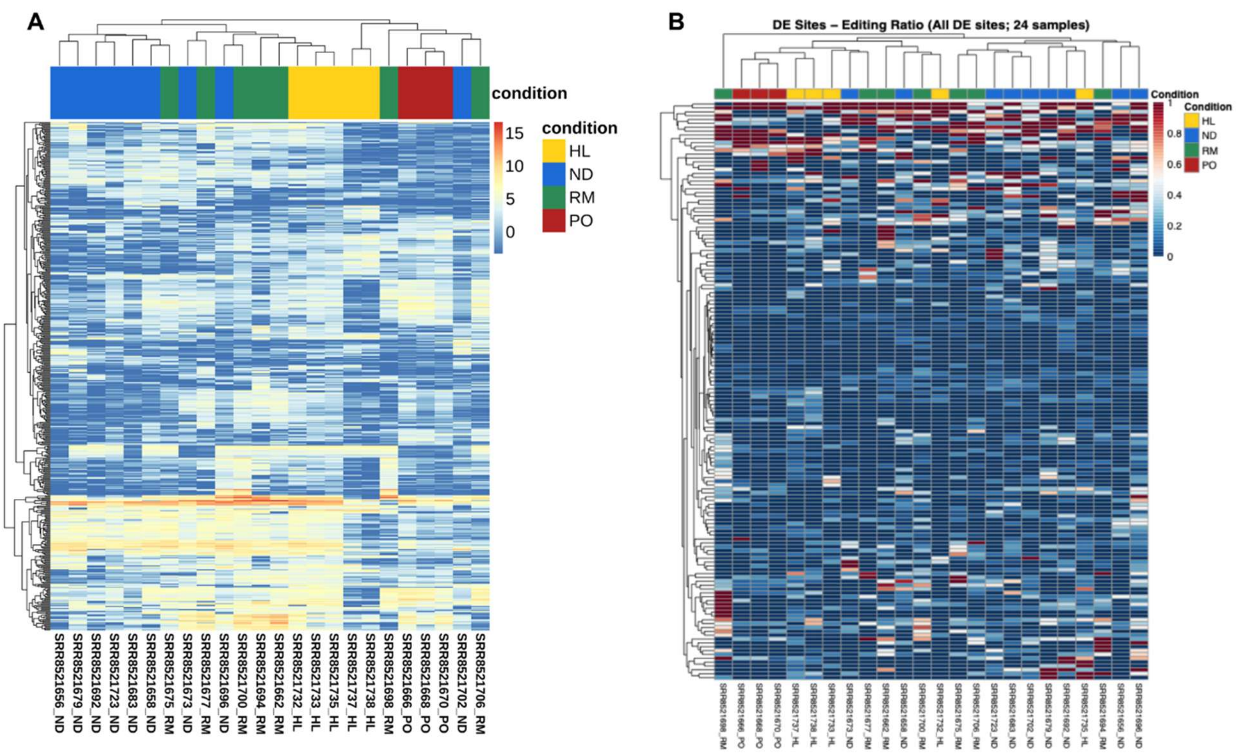


**Supplemental Figure 4. Heatmaps of gene expression and differential RNA editing across samples.** (A) Heatmap of the 500 most variable genes (rows) across samples (columns); values are $\log_2$(CPM). (B) Heatmap of differentially edited (DE) sites from the pseudo-bulk analysis (rows = DE sites, columns = samples); values are editing ratios. Column annotations denote clinical condition.
